# Host shifts result in parallel genetic changes when viruses evolve in closely related species

**DOI:** 10.1101/226175

**Authors:** Ben Longdon, Jonathan P Day, Joel M Alves, Sophia CL Smith, Thomas M Houslay, John E McGonigle, Lucia Tagliaferri, Francis M Jiggins

## Abstract

Host shifts, where a pathogen invades and establishes in a new host species, are a major source of emerging infectious diseases. They frequently occur between related host species and often rely on the pathogen evolving adaptations that increase their fitness in the novel host species. To investigate genetic changes in novel hosts, we experimentally evolved replicate lineages of an RNA virus (Drosophila C Virus) in 19 different species of Drosophilidae and deep sequenced the viral genomes. We found a strong pattern of parallel evolution, where viral lineages from the same host were genetically more similar to each other than to lineages from other host species. When we compared viruses that had evolved in different host species, we found that parallel genetic changes were more likely to occur if the two host species were closely related. This suggests that when a virus adapts to one host it might also become better adapted to closely related host species. This may explain in part why host shifts tend to occur between related species, and may mean that when a new pathogen appears in a given species, closely related species may become vulnerable to the new disease.

## Introduction

Host shifts where a pathogen jumps into and establishes in a new host species are a major source of emerging infectious diseases. RNA viruses seem particularly prone to host shift [1-4], with HIV, Ebola virus and SARS coronavirus all having been acquired by humans from other host species [5-7]. Whilst some pathogens may be pre-adapted to a novel host, there are increasing numbers of examples demonstrating that adaptation to the new host occurs following a host shift [8, 9]. These adaptations may allow a pathogen to enter host cells, increase replication rates, avoid or suppress the host immune response, or optimise virulence or transmission [10, 11]. For example, in the 2013-2016 Ebola virus epidemic in West Africa, a mutation in the viral glycoprotein gene that arose early in the outbreak and rose to high frequency was found to increase infectivity in human cells and decrease infectivity in bats, which are thought to be the source of Ebola virus [12, 13]. Likewise, a switch of a parvovirus from cats to dogs resulted in mutations in the virus capsid that allowed the virus to bind to cell receptors in dogs, but resulted in the virus losing its ability to infect cats [14, 15]

In some instances adaptation to a novel host relies on specific mutations that arise repeatedly whenever a pathogen switches to a given host. For example, in the jump of HIV-1 from chimps to humans, codon 30 of the *gag* gene has undergone a change that increases virus replication in humans, and this has occurred independently in all three HIV-1 lineages [5, 16]. Similarly, five parallel mutations have been observed in the two independent epidemics of SARS coronavirus following its jump from palm civets into humans [17]. Similar patterns have been seen in experimental evolution studies, where parallel genetic changes occur repeatedly when replicate viral lineages adapt to a new host species in the lab. For example, when Vesicular Stomatitis Virus was passaged in human or dog cells, the virus evolved parallel mutations when evolved on the same cell type [18]. Likewise, a study passaging Tobacco Etch Potyvirus on four plant species found parallel mutations occurred only when the virus infected the same host species [19]. These parallel mutations provide compelling evidence that these genetic changes are adaptive, with the same mutations evolving independently in response to natural selection [20].

The host phylogeny is important for determining a pathogens ability to infect a novel host, with pathogens tending to replicate most efficiently when they infect a novel host that is closely related to their original host [2, 21-34]. Here, we asked whether viruses acquire the same genetic changes when evolving in the same and closely related host species. We experimentally evolved replicate lineages of an RNA virus called Drosophila C Virus (DCV; Discistroviridae) in 19 species of Drosophilidae that vary in their relatedness and shared a common ancestor approximately 40 million years ago [35, 36]. We then sequenced the genomes of the evolved viral lineages and tested whether the same genetic changes arose when the virus was evolved in closely related host species.

## Results

### Parallel genetic changes occur in DCV lineages that have evolved in the same host species

To examine how viruses evolve in different host species we serially passaged DCV in 19 species of Drosophilidae. In total we infected 22,095 adult flies and generated 173 independent replicate lineages (6-10 per host species). We deep sequenced the evolved virus genomes to generate over 740,000 300bp sequence reads from each viral lineage. Out of 8989 sites, 584 contained a SNP with a derived allele frequency >0.05 in at least one viral lineage, and 84 of these were tri-allelic. None of these variants were found at an appreciable frequency in five sequencing libraries produced from the ancestral virus, indicating that they had spread though populations during the experiment (Figure 1). In multiple cases these variants had nearly reached fixation (Figure 1).

**Figure 1.**
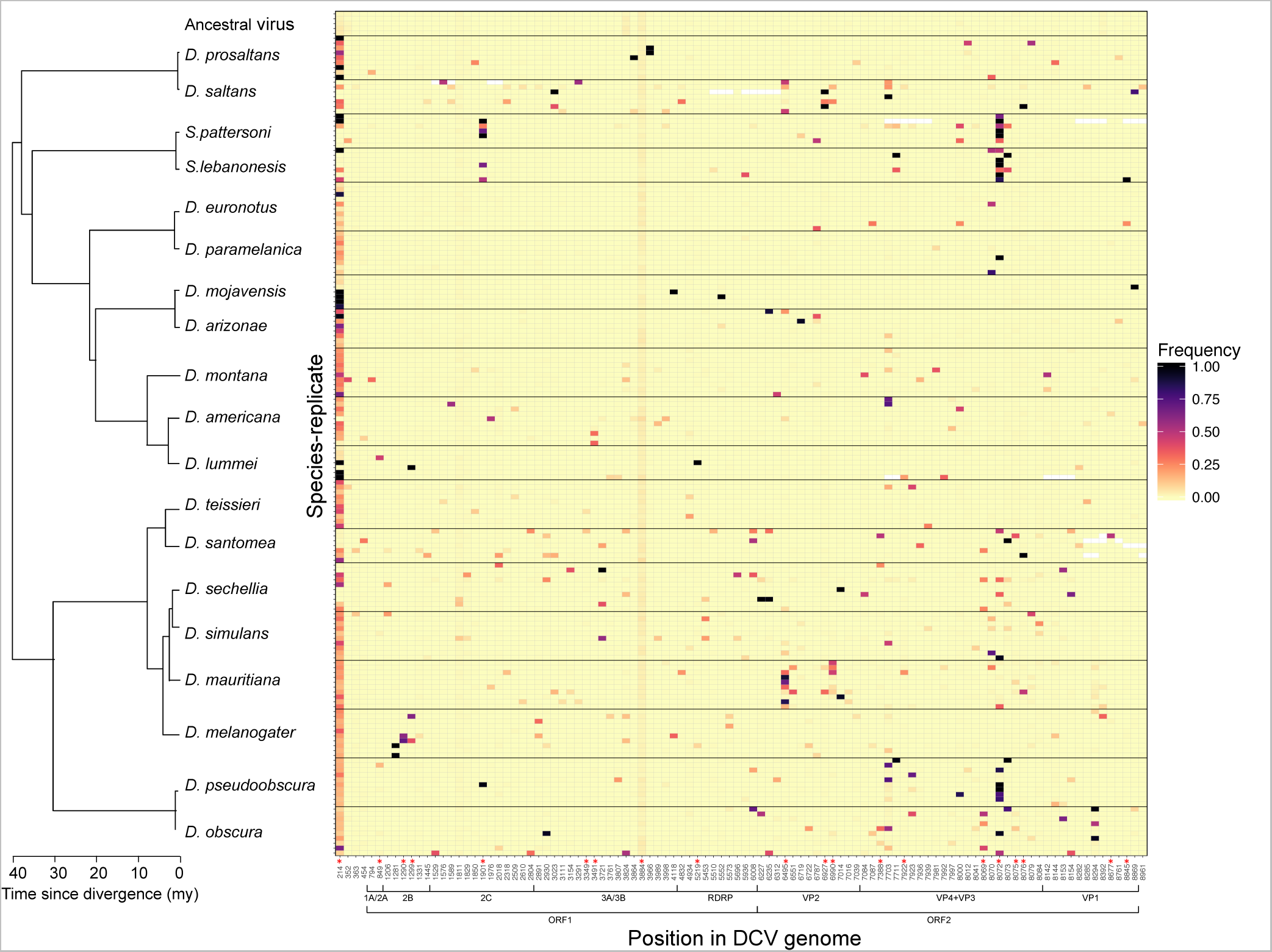
The frequency of SNPs in viral lineages that have evolved in different host species. Each row represents an independent viral lineage. Viruses that evolved in different host species are separated by black horizontal lines. Each column represents a polymorphic site in the DCV genome, and only sites where the derived allele frequency >0.05 in at least two lineages are shown. The intensity of shading represents the derived allele frequency. Sites where there are three alleles have the two derived allele frequencies pooled for illustrative purposes. Sites with SNP frequencies that are significantly correlated among lineages from the same host species are shown by red stars at the bottom the column (permutation test; *p*<0.05). Open reading frames (ORFs) and viral proteins based on predicted polyprotein cleavage sites [38-42] are below the x axis. Information on the distribution of mutations across the genome and whether they are synonymous or non-synonymous can be found in the supplementary results. Sites with missing data are shown in white. The phylogeny was inferred under a relaxed molecular clock [33, 43] and the scale axis represents the approximate age since divergence in millions of years (my) based on estimates from: [35, 36].

We next examined whether the same genetic changes occur in parallel when different populations encounterthe same host species. Of the 584 SNPs, 102 had derived allele frequencies >0.05 in at least two viral lineages, and some had risen to high frequencies in multiple lineages (Figure 1). We estimated the genetic differentiation between viral lineages by calculating *F*_ST_. We found that viral lineages that had evolved within the same host were genetically more similar to each other than to lineages from other host species (Figure 2; *P*<0.001). Furthermore, we found no evidence of differences in substitution biases in the different host species (Fisher Exact Test: *p*=0.14; see methods), suggesting that this pattern is not driven by changes in the types of mutations in different host species.

To examine the genetic basis of parallel evolution, we individually tested whether each SNP in the DCV genome showed a signature of parallel evolution among viral lineages passaged in the same host species (i.e. we repeated the analysis in Figures 2 for each SNP). We identified 56 polymorphic sites with a significant signal of parallel evolution within the same host species (*P*<0.05; significantly parallel sites are shown with a red asterisk in Figure 1; the false discovery rate is estimated to be 17% [37]).

**Figure 2.**
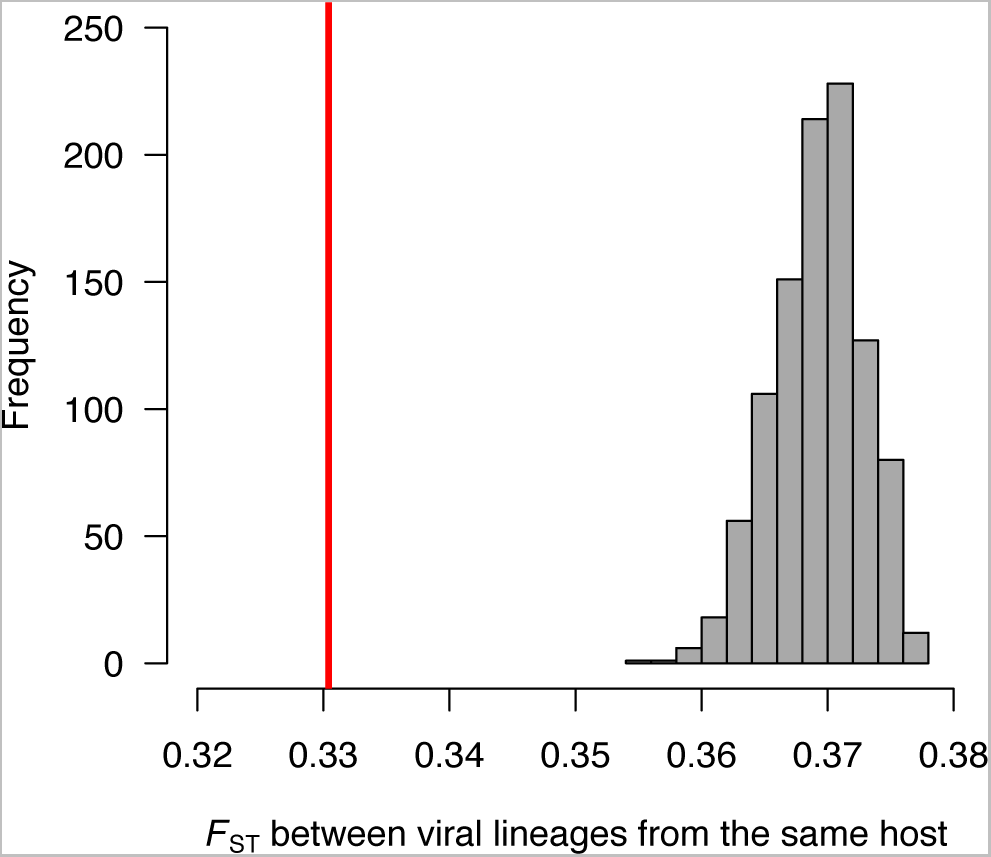
Viral lineages from the same host species were genetically more similar to each other than to lineages from different host species. The mean pairwise *F*_*ST*_ between all possible pairs of viral lineages from the same host species was calculated. The red line shows the observed value. The grey bars are the null distribution of this statistic obtained by permuting the viral lineages across host species 1000 times.

### Viruses in closely related hosts are genetically more similar

We investigated if viruses passaged through closely related hosts showed evidence of parallel genetic changes. We calculated *F*_ST_ between all possible pairs of viral lineages that had evolved in different host species. We found that viral lineages from closely related hosts were more similar to each other than viral lineages from more distantly related hosts (Figure 3A). This is reflected in a significant positive relationship between virus *F*_ST_ and host genetic distance (Figure 3B, Permutation test: *r*=0.15, *P*=0.002). We lacked the statistical power to identify the specific SNPs that are causing the signature of parallel evolution in Figure 3 (false discovery rate >0.49 for all SNPs).

**Figure 3.**
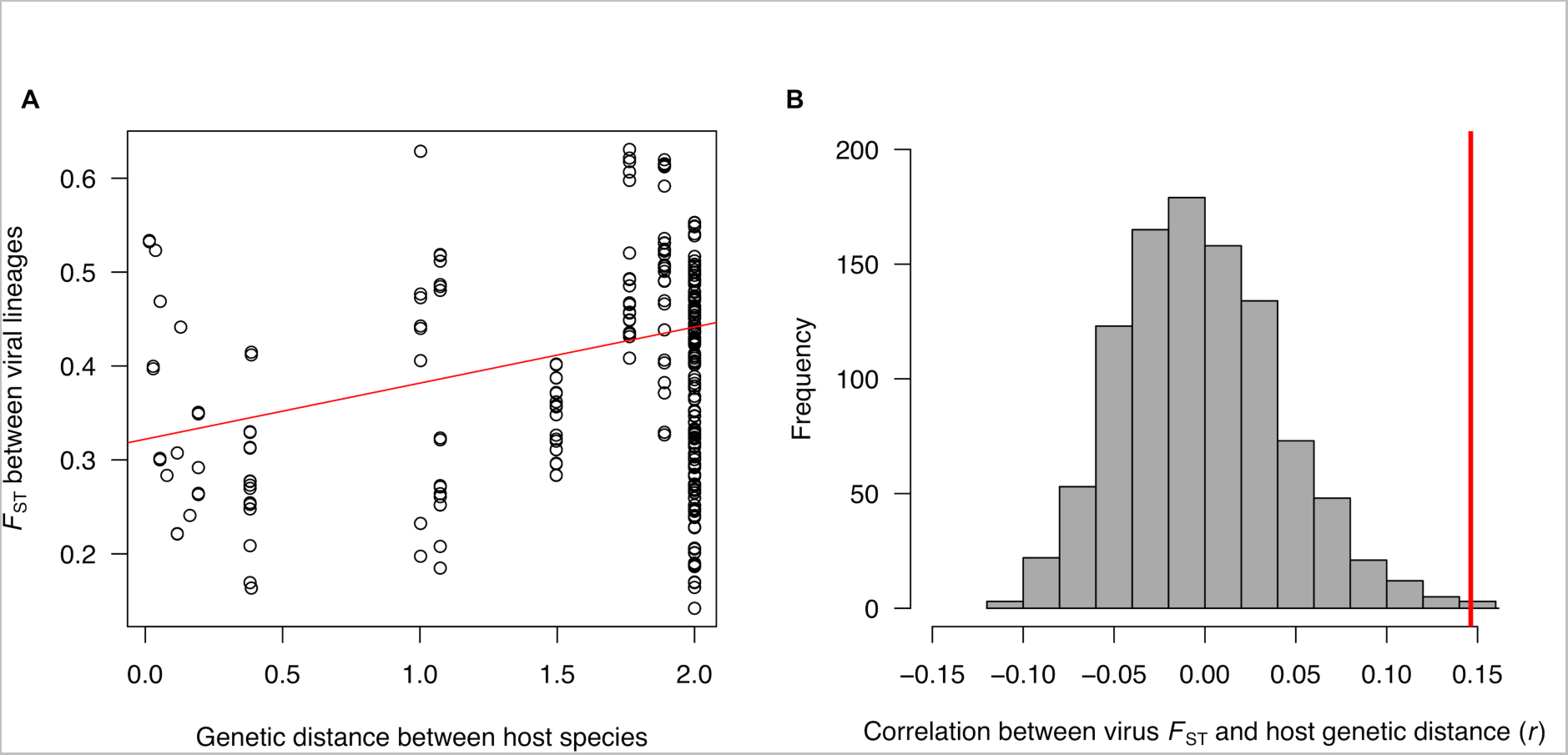
Viral lineages from more closely related host species are genetically more similar. (A) The correlation between the genetic differentiation of viral lineages and the genetic distance between the species they have evolved in. Linear regression line is shown in red. Genetic distances were scaled so that the distance from the root to the tip of the tree was one. (B) Pearson’s correlation coefficient (*r*) of *F*_*ST*_ between pairs of viral lineage and the genetic distance between the host species they evolved in. The observed value is in red and the grey bars are the null distribution obtained by permutation.

## Discussion

When a pathogen infects a novel host species, it finds itself in a new environment to which it must adapt [4, 8, 10, 44]. When DCV was passaged through different species of Drosophilidae, we found the same genetic changes arose repeatedly in replicate viral lineages in the same host species. Such repeatable parallel genetic changes to the same host environment are compelling evidence that these changes are adaptive [20]. We then examined whether these same genetic changes might occur in closely related host species, as these are likely to present a similar environment for the virus. We found that viruses evolved in closely related hosts were more similar to each other than viruses that evolved in more distantly related species. Therefore, mutations that evolve in one host species frequently arise when the virus infects closely related hosts. This finding of parallel genetic changes in closely related host species suggests that when a virus adapts to one host it might also become better adapted to closely related host species.

Phylogenetic patterns of host adaptation may in part explain why pathogens tend to be more likely to jump between closely related host species. This pattern is seen in nature, where host shifts tend to occur most frequently between closely related hosts, and in laboratory cross-infection studies, where viruses tend to replicate more rapidly when the new host is related to the pathogens natural host [2, 21-34]. For example, in a large cross-infection experiment involving Drosophila sigma viruses (Rhabdoviridae) isolated from different species of *Drosophila*, the viruses tended to replicate most efficiently in species closely related to their natural hosts [34]. This suggests that these viruses had acquired adaptations to their host species that benefitted them when they infected closely related species. Our results demonstrate that this pattern is apparent at the level of specific nucleotides, and can arise very shortly after a host shift. The function of these mutations is unknown, but in other systems adaptations after host shifts have been found to enhance the ability of the virus to bind to host receptors [11], increase replication rates [16] or avoid the host immune response [8, 10, 45].

While the susceptibility of a novel host is correlated to its relatedness to the pathogens’ original host, it is also common to find exceptions to this pattern. This is seen both in nature when pathogens shift between very distant hosts [46, 47], and in laboratory cross-infection experiments [33, 34]. This pattern is also seen in our data where we also observe parallel genetic changes occurring between more distantly related hosts. For example, a mutation at position 8072 was not only near fixation in most of the lineages infecting two closely related species, but also occurred at a high frequency in replicate lineages in a phylogenetically distant host (Figure 1).

In conclusion, we have found that host relatedness can be important in determining how viruses evolve when they find themselves in a new host. This study suggests that while some genetic changes will be found only in specific hosts, we frequently see the same changes occurring in closely related host species. These phylogenetic patterns suggest that mutations that adapt a virus to one host may also adapt it to closely related host species. Therefore, there may be a knock-on effect, where a host shift leaves closely related species vulnerable to the new disease.

## Methods

### Virus production

DCV is a positive sense RNA virus in the family Discistroviridae that was isolated from *D. melanogaster*, which it naturally infects in the wild [48, 49]. To minimise the amount of genetic variation in the DCV isolate we used to initiate the experimental evolution study, we aimed to isolate single infectious clones of DCV using a serial dilution procedure. DCV was produced in Schneider’s Drosophila line 2 (DL2) cells [50] as described in [51]. Cells were cultured at 25°C in Schneider’s Drosophila Medium with 10% Fetal Bovine Serum, 100 U/ml penicillin and 100 μg/ml streptomycin (all Invitrogen, UK). The DCV strain used was isolated from *D. melanogaster* collected in Charolles, France [52]. DL2 cells were seeded into two 96-well tissue culture plates at approximately 10^4^ cells in 100 μl of media per well. Cells were allowed to adhere to the plates by incubating at 25**°**C for five hours or over-night. Serial 1:1 dilutions of DCV were made in complete Schneider’s media, giving a range of final dilutions from 1:10^8^ 1:4x10^14^. 100 μl of these dilutions were then added to the cells and incubated for 7 days, 8 replicates were made for each DCV dilution. Each well was then examined for DCV infection of the DL2 cells, and a well was scored as positive for DCV infection if clear cytopathic effects were present in the majority of the cells. The media was taken from the wells with the greatest dilution factor that were scored as infected with DCV and stored at −80**°**C. This processes was then repeated using the DCV samples from the first dilution series. One clone, B6A, was selected for amplification and grown in cell culture as described above. Media containing DCV was removed and centrifuged at 3000 x g for 5 minutes at 4**°**C to pellet any remaining cell debris, before being aliquoted and stored at −80**°**C. The Tissue Culture Infective Dose 50 (TCID_50_) of the DCV was 6.32 x 10^9^ infectious particles per ml using the Reed-Muench end-point method [53].

### Inoculating fly species

We passaged the virus through 19 species of Drosophilidae, with 6-10 independent replicate passages for each species. We selected species from across the phylogeny (that shared a common ancestor approximately 40 million years ago [35, 36]), but included clades of closely related species that recently shared common ancestors less than 5 million years ago (Figure 1). All fly stocks were reared at 22**°**C. Stocks of each fly species were kept in 250ml bottles at staggered ages. Flies were collected and sexed, and males were placed on cornmeal medium for 4 days before inoculation. Details of the fly stocks used can be found in the supplementary materials.

4-11 day old males were infected with DCV using a 0.0125 mm diameter stainless steel needle (26002-10, Fine Science Tools, CA, USA) dipped in DCV solution. For the first passage this was the cloned DCV isolate in cell culture supernatant (described above), and then subsequently was the virus extracted from the previous passage (described below). The needle was pricked into the pleural suture on the thorax of flies, towards the midcoxa. Each replicate was infected using a new needle and strict general cleaning procedures were used to minimise any risk of cross-contamination between replicates. Species were collected and inoculated in a randomised order each passage. Flies were then placed into vials of cornmeal medium and kept at 22**°**C and 70% relative humidity. Flies were snap frozen in liquid nitrogen 3 days post-infection, homogenised in Ringer’s solution (2.5μl per fly) and then centrifuged at 12,000g for 10 mins at 4**°**C. The resulting supernatant was removed and frozen at −80**°**C to be used for infecting flies in the subsequent passage. The remaining homogenate was preserved in Trizol reagent (Invitrogen) and stored at −80°C for RNA extraction. The 3 day viral incubation period was chosen based on time course and pilot data showing that viral load reaches a maximum at approximately 3 days post-infection. This process was repeated for 10 passages for all species, except *D. montana* where only 8 passages were carried out due to the fly stocks failing to reproduce. Each lineage was injected into a mean of 11 flies at each passage (range 4-18). Experimental evolution studies in different tissue types have seen clear signals of adaptation in 100 virus generations [18]. Based on log_2_ change in RNA viral load we estimate that we have passaged DCV for approximately 100-200 generations.

### Sequencing

After passaging the virus, we sequenced evolved viral lineages from 19 host species, with a mean of 9 independent replicate lineages of the virus per species (range 6-10 replicates). cDNA was synthesised using Invitrogen Superscript III reverse-transcriptase with random hexamer primers (25**°**C 5mins, 50**°**C 50mins, 70**°**C 15mins). The genome of the evolved viruses, along with the initial DCV ancestor (x5) were then amplified using Q5 high fidelity polymerase (NEB) in nine overlapping PCR reactions (see supplementary Table S2 for PCR primers and cycle conditions). Primers covered position 62-9050bp (8989bp) of the Genbank refseq (NC_001834.1) giving 97% coverage of the genome. PCRs of individual genomes were pooled and purified with Ampure XP beads (Agencourt). Individual Nextera XT libraries (Illumina) were prepared for each viral lineage. In total we sequenced 173 DCV pooled amplicon libraries on an Illumina MiSeq (Cambridge Genomic Service) v3 for 600 cycles to give 300bp paired-end reads.

### Bioinformatics and variant calling

FastQC, version 0.11.2 [54] was used to assess read quality and primer contamination. Trimmomatic, version 0.32 [55] was used to removed low quality bases and adaptor sequences, using the following options: MINLEN=30 (Drop the read if it is below 30 base pairs), TRAILING=15 (cut bases of the end of the read if below a threshold quality of 15), SLIDINGWINDOW=4:20 (perform a sliding window trimming, cutting once the average quality within a 4bp window falls below a threshold of 20), and ILLUMINACLIP=TruSeq3-PE.fa:2:20:10:1:true (remove adapter contamination; the values correspond in order to: input fasta file with adapter sequences to be matched, seed mismatches, palindrome clip threshold, simple clip threshold, minimum adapter length and logical value to keep both reads in case of read-through being detected in paired reads by palindrome mode).

To generate a reference ancestral Drosophila C Virus sequence we amplified the ancestral starting virus by PCR as above. PCR products were treated with exonuclease 1 and Antarctic phosphatase to remove unused PCR primers and dNTPs and then sequenced directly using BigDye reagents (ABI) on an ABI 3730 capillary sequencer in both directions (Source Bioscience, Cambridge, UK). Sequences were edited in Sequencher (version 4.8; Gene Codes), and were manually checked for errors. Fastq reads were independently aligned to this reference sequence (Genbank accession: MG570143) using BWA-MEM, version 0.7.10 {Li, 2009 #1605} with default options with exception of the parameter–M, which marks shorter split hits as secondary. 99.5% of reads had mapping phred quality scores of >60. The generated SAM files were converted to their binary format (BAM) and sorted by their leftmost coordinates with SAMtools, version 0.1.19 (website: http://samtools.sourceforge.net/) [56]. Read Group information (RG) was added to the BAM files using the module AddOrReplaceReadGroups from Picard Tools, version 1.126 (https://broadinstitute.github.io/picard).

The variant calling was then performed for each individual BAM using UnifiedGenotyper tool from GATK, version 3.3.0. As we were interested in calling low frequency variants in our viruses, we assumed a ploidy level of 100 (-sample_ploidy:100). The other parameters were set to their defaults except-stand_call_conf:30 (minimum phred-scaled confidence threshold at which variants should be called) and-downsample_to_coverage:1000 (down-sample each sample to 1000X coverage)

### Host phylogeny

We used a trimmed version of a phylogeny produced previously [33]. This time-based tree (where the distance from the root to the tip is equal for all taxa) was inferred using seven genes with a relaxed molecular clock model in BEAST (v1.8.0) [43, 57]. The tree was pruned to the 19 species used using the Ape package in R [58, 59].

### Statistical Analysis

We examined the frequency of alternate alleles (single nucleotide polymorphisms: SNPs) in five ancestral virus replicates (aliquots of the same virus stock that was used to found the evolved lineages). SNPs in these ancestral viruses may represent pre-standing genetic variation, or may be sequencing errors. We found the mean SNP frequency was 0.000923 and the highest frequency of any SNP was 0.043 across the ancestral viruses. We therefore included a SNP in our analyses if its frequency was >0.05 in any of the evolved viral lineages. For all analyses we included all three alleles at triallelic sites.

### Parallel evolution within species

As a measure of genetic differentiation we estimated *F*_*ST*_ between all the virus lineages based on the heterozygosity (*H*) of the SNPs we called [60]:

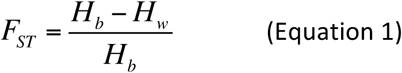

where *H*_*b*_ is the mean number of differences between pairs of sequence reads sampled from the two different lineages. *H*_*w*_ is mean number of differences between sequence reads sampled from within each lineage. *H*_*b*_ and *H*_*w*_ were calculated separately for each polymorphic site, and the mean across sites used in equation (1). *H*_*w*_ was calculated separately for the two lineages being compared, and the unweighted mean used in equation (1).

To examine whether there had been parallel evolution among viral lineages that had evolved within the same fly species, we calculated the mean *F*_*ST*_ between lineages that had evolved in the same fly species, and compared this to the mean *F*_*ST*_ between lineages that had evolved in different fly species. We tested whether this difference was statistically significant using a permutation test. The fly species labels were randomly reassigned to the viral lineages, and we calculated the mean *F*_*ST*_ between lineages that had evolved in the same fly species. This was repeated 1000 times to generate a null distribution of the test statistic, and this was then compared to the observed value.

To identify individual SNPs with a signature of parallel evolution within species, we repeated this procedure separately for each SNP.

### Parallel evolution between species

We next examined whether viral lineages that had evolved in different fly species tended to be more similar if the fly species were more closely related. Considering all pairs of viral lineages from different host species, we correlated pairwise *F*_*ST*_ with the genetic distance between the fly species. To test the significance of this correlation, we permuted the fly species over the *Drosophila* phylogeny and recalculated the Pearson correlation coefficient. This was repeated 1000 times to generate a null distribution of the test statistic, and this was then compared to the observed value. To identify individual SNPs whose frequencies were correlated with the genetic distance between hosts we repeated this procedure separately for each SNP.

We confirmed there was no relationship between rates of molecular evolution (SNP frequency) and either genetic distance from the host DCV was isolated from (*D. melanogaster*) or estimated viral population size (see supplementary Figures S1 and S2) using generalised linear mixed models that include the phylogeny as a random effect in the MCMCglmm package in R [61] as described previously [34]. We also examined the distribution of SNPs and whether they were synonymous or non-synonymous (see supplementary results).

To test whether there were systematic differences in the types of mutations occurring in the different host species, we classified all the SNPs into the six possible types (A/G, A/T, A/C, G/T, G/C and C/T). We then counted the number of times each type of SNP arose in each host species at a frequency above 5% and in at least one biological replicate (SNPs in multiple biological replicates were only counted once). This resulted in a contingency table with 6 columns and 19 rows. We tested for differences between the species in the relative frequency of the 6 SNP types by simulation [62].

Sequence data (fastq files) are available in the NCBI SRA (Accession: SRP119720). BAM files, data and R scripts for analysis in the main text are available from the NERC data repository (funding requirement - awaiting doi, temporary link to data and scripts https://figshare.com/s/b119ba86def8bca58782).

## Author contributions

Designing experiment: BL, FMJ. Lab work: JPD, SCLS, TMH, LT, BL Bioinformatics analysis: JMA, JEM, FMJ, BL. Statistical analysis: BL, FMJ. Manuscript written by BL and FMJ with input from all authors.

## Acknowledgements

Thanks to the Drosophila species stock centre for providing fly stocks and four anonymous reviewers for constructive comments.

